# New genotype invasion of dengue virus serotype 1 drove massive outbreak in Guangzhou, China

**DOI:** 10.1101/697052

**Authors:** Qinlong Jing, Sean Wu, Zhengjian He, Lihong Yuan, Mengmeng Ma, Zhijun Bai, Liyun Jiang, John Marshall, Jiahai Lu, Zhicong Yang

## Abstract

**Background:** Dengue fever is a mosquito-borne infectious disease that has caused major health problems. Variations in dengue virus (DENV) genes are important features of epidemic outbreaks. However, the associations of DENV genes with epidemic scale have not been extensively examined. Here, we assessed new genotype invasion of DENV-1 isolated from Guangzhou in China to evaluate associations with epidemic outbreaks.

**Methodology/Principal Findings:** We used DENV-1 strains isolated from sera of dengue cases from 2002 to 2016 in Guangzhou for complete genome sequencing. A neighbor-joining phylogenetic tree was constructed to elucidate the genotype characteristics and determine if new genotype invasion correlated with major outbreaks. In our study, a new genotype invasion event was observed during each significant outbreak period in 2002-2003, 2006-2007 and 2013-2014. Genotype II was the main epidemic genotype in 2003 and before. Invasion of genotype I in 2006 caused an unusual outbreak with 765 cases (relative risk (RR)=16.24, 95% confidence interval (CI) =12.41-21.25). At the middle and late stages of the 2013 outbreak, genotype III was introduced to Guangzhou as a new genotype invasion responsible for 37340 cases with RR 541.73 (95%CI=417.78-702.45), after which genotypes I and III began co-circulating. Base mutations occurred after new genotype invasion, and the gene sequence of NS3 protein had the lowest average similarity ratio (99.82%), followed by the gene sequence of E protein (99.86%), as compared to the 2013 strain.

**Conclusions/Significance:** Genotype replacement and co-circulation of multiple DENV-1 genotypes were observed. New genotype invasion was highly correlated with local unusual outbreaks. In addition to DENV-1 genotype I in the unprecedented outbreak in 2014, new genotype invasion by DENV-1 genotype III occurred in Guangzhou.

**Author Summary:** New genotype invasion of dengue virus highly correlates with the massive outbreaks. In this study, we examined the association of the genotype of dengue virus serorype 1 (DENV-1) from human cases through complete genome sequencing with outbreak scale during 2002 and 2016 in Guangzhou, China. It was observed that genotype replacement and co-circulation of multiple genotypes occurred. Most importantly, it indicated that new genotype invasion was highly related with local unusual outbreaks in major outbreak periods in 2002-2003, 2006-2007 and 2013-2014. DENV-1 genotype II was the main epidemic genotype in 2003 and before. Invasion of genotype I in 2006 caused an unusual outbreak with 765 cases reported. In addition to genotype I circulation, new genotype invasion by genotype III was the key determinant for the 2014 massive outbreak reaching the highest number of cases with 37340. Furthermore, base mutations appeared after genotype III invasion, and the gene sequence of NS3 protein had the lowest average similarity ratio, followed by the gene sequence of E protein, as compared to the 2013 strain.

## Introduction

Dengue fever (DF), transmitted by the bite of infected *Aedes* mosquitoes, has become the most rapidly spreading arboviral disease in recent decades, with accelerating expansion in affected geographic regions worldwide. Currently, approximately half of the world’s population lives in areas at risk of infection. Three hundred ninety million infections and 96 million symptomatic cases occur annually, among which 500,000 individuals suffer from severe dengue, such as dengue hemorrhagic fever or dengue shock syndrome [1, 2]. The disease has huge health and economic effects, with an estimated burden of 25.5 disability-adjusted life years per 100,000 individuals [3]. However, application of the only currently licensed vaccine, CYD-TDV (Dengvaxia; Sanofi Pasteur, Lyon, France) is limited owing to safety issues associated with increased hospitalization risk for individuals who have never been infected with dengue before [4], and no specific interventions to treat the disease have been established.

Dengue virus (DENV), a single-stranded positive-sense RNA virus with an 11-kb genome, contains an open reading frame encoding three structural proteins (C, prM/M, and E) and seven nonstructural proteins (NS1, NS2A, NS2B, NS3, NS4A, NS4B, and NS5). The virus can be further classified into four serotypes according to their distinctive antigenicities, i.e., DENV-1–4, and there are diverse genotypes within each serotype. The sequence of each DENV genotype is not always fixed, and frequent variations, recombinations, and lineage turnover or replacement may occur because of selection pressure [5].

In recent decades, DF has become a major threat to public health in Southern China [6]. Frequent and large-scale outbreaks have posed a huge disease burden; in particular, serious outbreaks dominated by DENV-1 have occurred in the last 20 years. Guangzhou has become the most heavily affected area in China since the 1990s; the number of reported cases during 2005–2017 accounted for 60.86% of all cases nationally, reaching 81% in 2014 alone [7–9]. Outbreaks with highest incidence rates were all caused by DENV-1, leading to extensive studies of associated factors, such as climate, mosquito density, control measures, and virus serotypes [10–13]. Despite this, numerous scientific issues related to large-scale outbreaks remain unresolved, particularly the cause of the massive outbreak in Guangzhou in 2013 and 2014.

Genetic variations in DENV are important factors contributing to the severity of epidemics. However, the associations of such genetic variations with epidemic scale have not been thoroughly examined [2, 14]. Virus variations and resulting invasion by new genotypes may be key factors driving large-scale outbreaks and the development of severe symptoms and death when other confounding factors, such as changes in mosquito vectors, climate, imported cases, tourism, and trade, are not altered [15]. Previous studies have mostly focused on E gene and prM/M gene fragments to analyze genetic variations in DENV, and few studies have reported analysis of complete genome sequences. Analysis based on the E gene indicated that DENV genes exhibit diverse lineages and geographical distributions. Different serotypes comprise various subgroups called genotypes, which can differ in virus virulence and transmission rate [16]. DENV-1 has five different genotypes (I–V), among which genotypes I, IV, and V are still prevalent, whereas genotypes II and III appear to have become dormant. Phylogenetic analyses have shown that different virus isolates with different lineage features nonetheless clustered within the same genotype [17]. The practicality and development of high-throughput whole-genome sequencing and deep sequencing have enabled application of new analytical methods. A recent complete genome sequence analysis revealed that DENV-1 could be classified into three genotypes (I, II, and III), in contrast to the results of previous genotyping based on the E gene [18]. Lee et al. also reported that DENV-1 could be classified into three genotypes by complete genome sequencing [19] in an investigation of a historically large-scale outbreak in Singapore during 2013 and 2014. Their findings showed that the outbreak was related to the introduction of a new genotype III; however, the roles of different genotypes in driving outbreaks have not been sufficiently evaluated.

There are three scenarios under which new genotype invasion may occur in a specific area. First, genetic variation worldwide produces new genotypes. Second, genotypes that are prevalent in local areas are transmitted to areas in which there are no such genotypes. Third, genotypes that have been silent for years suddenly emerge. The status of dengue in China is still that of an imported disease, that can trigger local transmissions [20]. Guangzhou has emerged as an important hotspot, with the number of reported cases exceeding half those recorded nationwide. Currently, Guangzhou is still regarded as a nonendemic area for dengue, supported by numerous studies of mathematical and statistical modeling, virus evolution, and epidemiology [10, 19, 20]. In mainland China, particularly Guangzhou, many studies have evaluated serotyping and genotyping based on the E gene. However, sequence analysis based on the complete genome has not been performed deeply, and efforts to elucidate the impact of introduction of new genotypes on outbreaks have been insufficient.

Because DENV-1 is a representative serotype causing dengue outbreaks in Guangzhou, in this study, we analyzed new genotype invasion and the capacity of different genotypes to drive outbreaks by phylogenetic analysis based on DENV-1 complete genome sequences. We also explored the hypothesis of new genotype invasion as the main driver in major outbreak years. Our findings are expected to provide insights into viral evolutionary dynamics and the potential causes of massive outbreaks, which will help improve prevention and control measures for dengue.

## Methods

### Data sources and case investigations

The 2001–2005 case data were obtained from the archives of the Guangzhou Center for Disease Control and Prevention (GZCDC), including case questionnaires, epidemiological survey reports, phase analysis reports, and summaries. The 2006–2016 case information was extracted from the National Notifiable Infectious Diseases Reporting Information System of China. Once medical institutes reported suspected cases through the system, the local district CDC staff would conduct face-to-face case investigations to collect data on demographics, disease information, clinical manifestations, and travel history (domestic and international). Serum samples were also collected at this time.

### Specimen collection

DENV strains were isolated from serum specimens of reported DF cases and preserved by the GZCDC and Sun Yat-sen University (SYSU). Blood samples (3–5 mL) from the acute phase (within 6 days after the date of onset) were collected from consenting patients by a nurse at the visiting medical institute or field CDC staff and then separated to obtain serum. Sera were stored at −80°C until processing.

### Serotyping and virus isolation

We followed serotyping and virus isolation assay protocols recommended by the World Health Organization [21]. Briefly, all serum samples were extracted with total viral RNA using a QIAamp viral RNA mini kit (Qiagen, Germany). Next, a TaqMan probe-based real-time polymerase chain reaction (PCR) protocol was employed to determine the serotype. Positive sera were diluted 10-fold with the sample treatment solution at 4°C for 2 h before being inoculated onto an *Aedes albopictus* mosquito (C6/36) cell line for virus isolation. Cultures were passaged no more than three times. The experiments were conducted in laboratories at GZCDC and by our collaborator SYSU in Guangzhou.

### DENV complete genome sequencing

We selected representative virus strains and variants for complete genome sequencing. Specific primers for amplification and sequencing are listed in Additional File 1 [22], and primer synthesis was performed by a qualified third-party biotech company. The PCR product was purified using a QIAquick gel extraction kit (Qiagen) according to the manufacturer’s instructions and then sent to a qualified third-party biotech company for sequencing using an ABI 3730xl DNA analyzer platform (Applied Biosystems, CA, USA). Nucleotide sequences were initially assembled with Lasergene software (version 8.0; DNASTAR Inc., Madison, WI, USA), and continuous sequences were aligned using BioEdit software (version 7.0.5). In addition, selected strains were transferred to a third-party biotech company (BGI, China) for complete genome sequencing by next-generation sequencing.

### Phylogenetic analysis

DENV-1 reference strains were downloaded from GenBank, and those with a length greater than 10000 bp and associated metadata containing the year and location the isolate was sampled were included for phylogenetic analysis. Included samples were from different countries and regions worldwide, and the isolation year varied from 1944 to 2016. During complete genome sequence analysis, sequences identified from the same country in the same year with an evolutionary distance of zero were excluded through multiple sequence alignment.

All complete genome sequences were subjected to multiple sequence alignment using Mafft version 7 (https://mafft.cbrc.jp/alignment/software/). We then performed sequence similarity analysis with Mega 7.0 software (https://www.megasoftware.net/) and constructed neighbor-joining phylogenetic trees using Kimura’s two-parameter model. The robustness of nodes was assessed with 1000 bootstrap replicates. All reference strains were labeled with name, year of isolation, location of isolation, and GenBank accession number. For strains other than Chinese strains, only one reference strain was randomly selected for phylogenetic analysis when multiple strains were isolated from the same year and the same country and when their evolutionary distance was zero after multiple sequence alignment by the K80 model.

### Association of new genotype invasion with epidemic scale in major outbreak years

In this study, outbreak year was defined as the year when the annual number of local cases exceeded 500. From 2001 onwards, major outbreak years in Guangzhou have often appeared in two consecutive years, including 2002 and 2003, 2006 and 2007, and 2013 and 2014. Therefore, these three periods were considered three distinct outbreak years in the analysis. Here, we investigated the association of new genotype invasion with incidence rate by including new genotype as the study factor and the incidence rate in each outbreak year as the dependent variable, while using the average incidence rate during study years (median incidence rate) as the control. The incidence rates were compared with relative risks (RRs) and 95% confidence intervals (95% CIs) calculated to determine whether the effects of genotype were statistically significant.

### Comparison of the capacity of different genotypes to drive DF outbreaks during the same epidemic period

Because 2013 and 2014 were the most important years for DF outbreaks in Guangzhou, we evaluated the ability of genotype III to drive more severe infections in comparable communities during these two periods. Comparable communities that shared genotype III or genotype I outbreaks were selected based on similar sizes of permanent resident populations and local environment types. We then calculated RRs and 95% CIs to determine the capacity of different genotypes to drive the outbreak during the same epidemic period.

### Association of different genotypes with outbreaks during 2002–2016

In order to explore differences in the capacities of various genotypes to drive outbreaks in different years, we employed univariate and multivariate linear regression models. Community incidence caused by different genotypes was used as the dependent variable, genotype was used as the independent variable, and population density was used as the adjusting variable. The models were tested by analysis of variance with regression coefficients confirmed by *t* tests, and factors with *P* values of less than 0.05 were retained in the final model.

### Statistical analysis and graphing

We used R (version 3.2.2; the R Foundation for Statistical Computing, Vienna, Austria) to process demographic, epidemiological, and genomic data using dplyr and ape packages. The geographic source and other general information for DENV strains were evaluated using descriptive analyses.

### Ethics statement

The research protocol was reviewed and approved by the institutional review boards of both the GZCDC and the School of Public Health, SYSU. Written informed consent was obtained from adult participants (age ≥ 18 years old) or parents or legal guardians of children enrolled in the study (age < 18 years old). Consent was also obtained from children ages 7–18 years old.

## Results

In total, 1679 DENV-1 complete genome sequences were included in the phylogenetic analysis, including 97 strains from China and 1582 strains from other countries and regions, such as Southeast Asia, East Asia, South America, Central America, Africa, the Middle East, and Europe (Additional File Table S2).

As shown in Table 1, there were 65 DENV-1 strains identified from Guangzhou since 1991, accounting for 67.01% (65/97) of all Chinese strains included in the analysis, among which 48 (73.85%, 48/65) strains were sequenced from 204 serum specimens by our research team and collaborators.

**Table 1.** DENV-1 genome sequences in Guangzhou in different years

| Year | Number | Proportion (%) |
| --- | --- | --- |
| 1991 | 1 | 1.54 |
| **1995** | **1** | **1.54** |
| 1999 | 1 | 1.54 |
| **2002** | **3** | **4.62** |
| 2003 | 2 | 3.08 |
| **2006** | **6** | **9.23** |
| 2007 | 3 | 4.62 |
| 2010 | 1 | 1.54 |
| 2011 | 1 | 1.54 |
| **2013** | **8** | **12.31** |
| **2014** | **30** | **46.15** |
| 2015 | 2 | 3.08 |
| 2016 | 6 | 9.23 |
| Total | 65 | 100.00 |

### DENV-1 genome genotyping

Phylogenetic analysis using complete genome sequences showed that DENV-1 was generally classified into three genotypes, i.e., genotype I, II, and III (Additional File Figure S1). All three genotypes have been observed in DENV-1 outbreaks in China, and genotype III was believed to be new in China at the time of its detection. Specifically, DENV-1 genotype III was first identified during the large outbreak in 2013–2014 in Guangzhou, demonstrating highest similarity with strains from India (JQ922548/India/2005, JQ917404/India/2009) and Singapore (KM403584/Singapore/2013), and no prior outbreaks of genotype III had been recorded in China.

### DENV-1 genome genotyping of Guangzhou isolates

As shown in complete genome sequence analysis of 65 DENV-1 strains from Guangzhou, genotypes I, II, and III have all been found circulating in Guangzhou. Large-scale outbreaks typically occurred in the years when a genotype was introduced for the first time, such as 2002, 2006, 2013, and 2014 (Figures 1 and 2).

**Fig 1.**
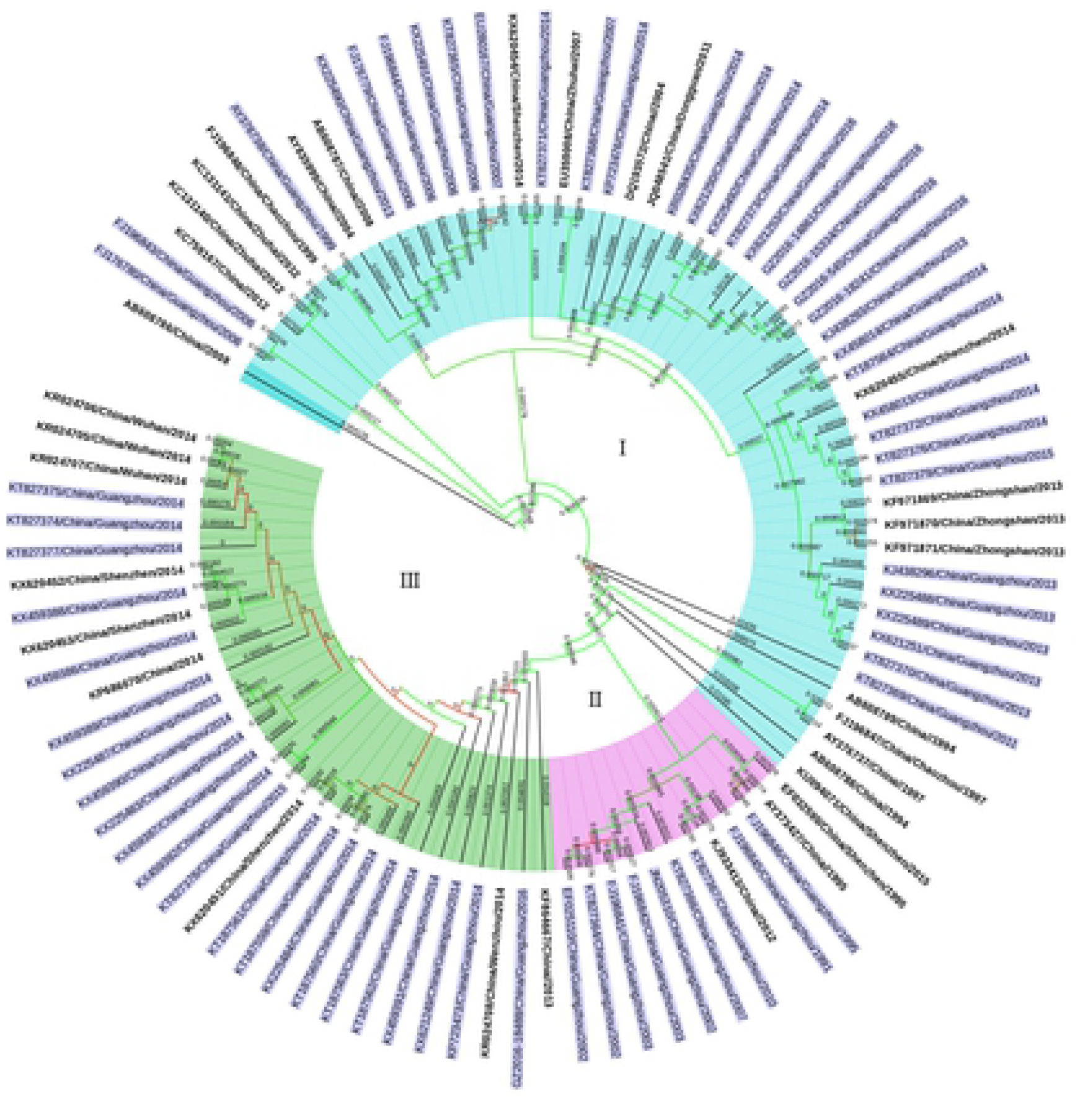
Phylogenetic tree of complete genome sequences of Chinese DENV-1 strains. Guangzhou strains are labeled in light blue. Strains of genotypes I, II, and III are highlighted in light cyan, lavender, and light green, respectively.

**Fig 2.**
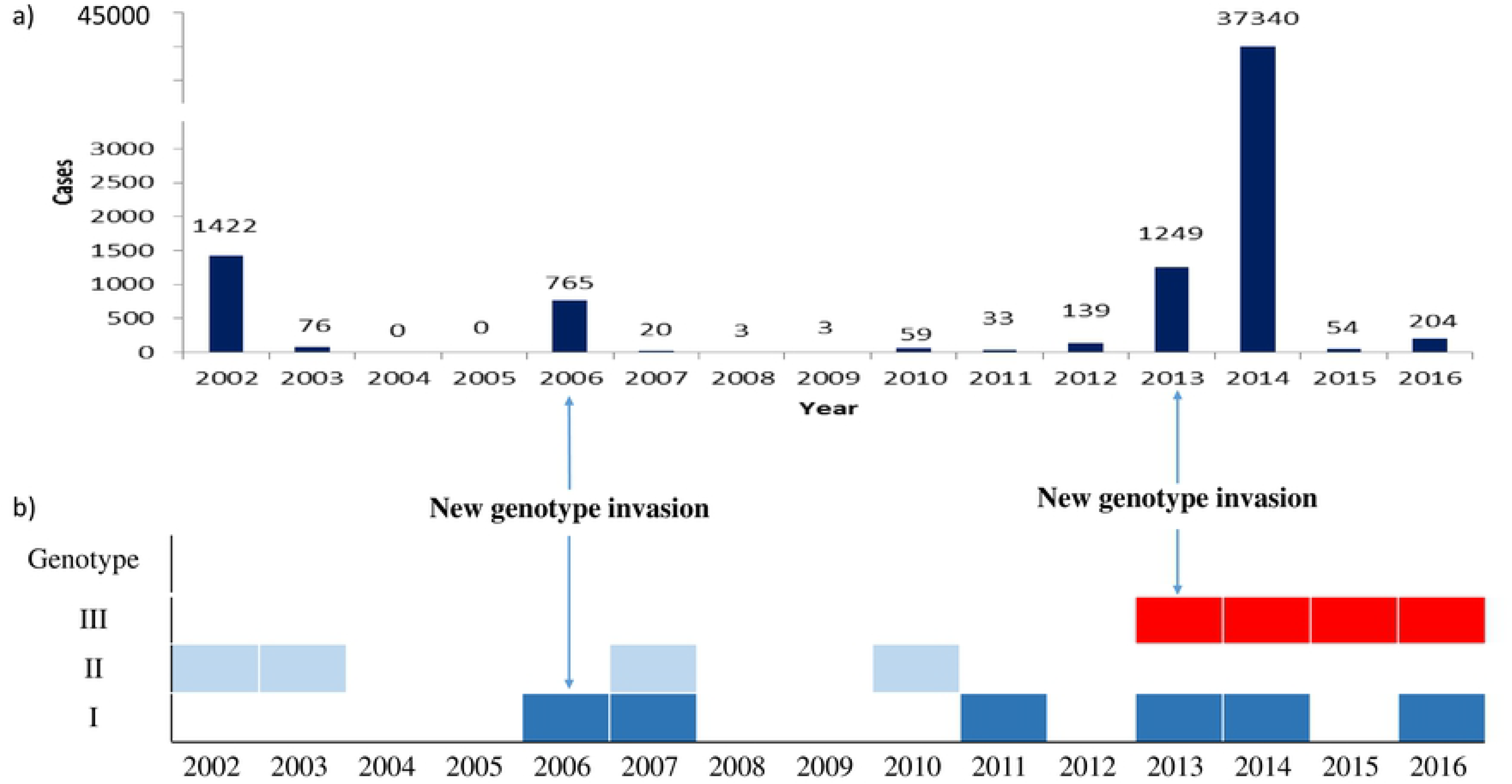
Numbers of local dengue cases and yearly genotype distributions from 2002 to 2016. a) Number of local cases DF reported in Guangzhou (2002–2016). b) Yearly distribution of DENV-1 genotypes in Guangzhou (2002–2016). The year is indicated when a new genotype invasion was introduced.

In 2006, DENV-1 genotype I was introduced into Guangzhou as a new genotype invasion, causing a large-scale outbreak in that year; 765 local cases were reported, followed by a recurrent outbreak of 20 cases in 2007. During the following years until 2012, no genotype I outbreaks were observed except for a small-scale outbreak in 2011 (33 cases). However, this genotype re-emerged in 2013, 2014, and 2016.

DENV-1 genotype II was first isolated in Guangzhou in 1991 and triggered the largest epidemic to date in 1995, with 5337 local cases reported. Later outbreaks occurred in 2002 (1422 cases) and 2003 (76 cases). However, there were no large-scale genotype II outbreaks in Guangzhou (76 cases in 2003, 20 cases in 2007, and 59 cases in 2010) after 2002.

DENV-1 genotype III was first introduced into Guangzhou in the form of a new genotype invasion in 2013. In that year, genotypes I and III co-circulated, and 1249 local cases were reported. Subsequently, in 2014, a historically unprecedented epidemic of local cases was observed in Guangzhou, with a total of 37,340 local cases reported. In April 2015, genotype III was isolated again in Guangzhou. Nevertheless, there were no further DENV-1 cases after June 2015 until 2016. Overall, DENV-2 was mainly responsible for the DF outbreak in Guangzhou in 2015.

### Association of new genotype invasion with epidemic scale in major outbreak years

The incidence rates triggered by different genotypes of new genotype invasion in different years were all higher than the average rates during the study years (median number of reported local cases from 2001 to 2016). Because our viral isolation procedures were initiated in 2002, we included 2002 as the year of invasion of genotype II. We found that genotype III showed the greatest capacity for driving outbreaks, with an RR value as high as 541.73 (95% CI: 417.78–702.45), followed by genotype II, with a RR value of 37.83 (95% CI: 29.02–49.26). The capacity of genotype I was relatively lower. However, during the specific year of invasion (2006–2007), the RR value of genotype I still reached 16.24 (95% CI: 12.41–21.25). In general, the epidemic capacity of new genotype invasion in Guangzhou varied, with genotype III being the strongest, and the risk was 14.32 and 33.36 times higher than those of genotypes I and II, respectively (Table 2).

**Table 2.**
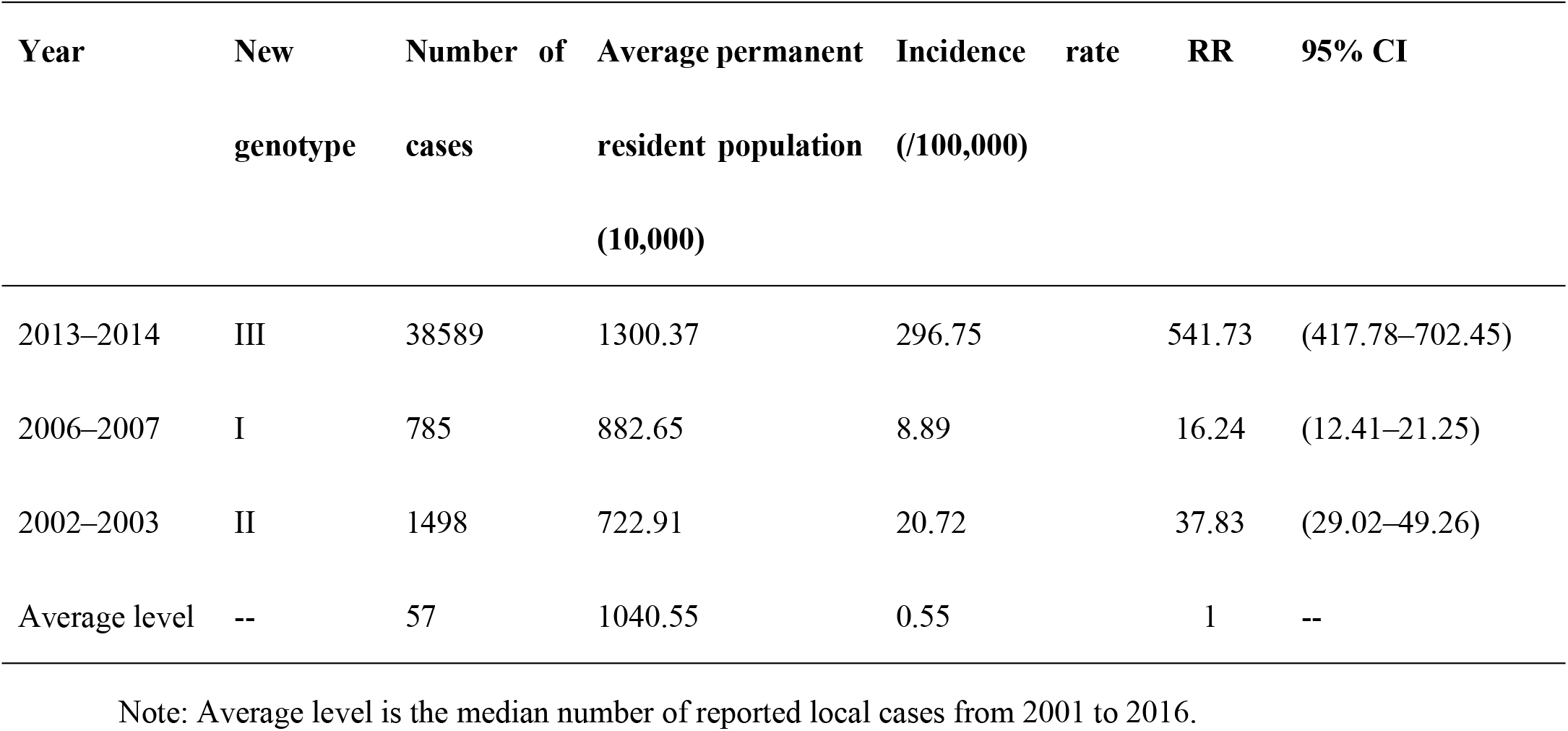
Comparison of new genotype invasion and epidemic scale during different years

### Comparison of the capacities of different genotypes for driving outbreaks during the same epidemic period

Among 48 DENV-1 strains sequenced by our research team, there were 36 (75%) strains isolated from different DF cases with clear background information, including the patient’s permanent address in Guangzhou and time of onset of symptoms. These strains included 14 strains of genotype I, three strains of genotype II, and 19 strains of genotype III across various years, with one in 2002, one in 2006, three in 2007, two in 2010, one in 2011, five in 2013, 17 in 2014, two in 2015, and four in 2016. Because of the extremely large outbreak observed during 2013–2014, this period was selected as the study epidemic period to compare the capacities of different genotypes for driving DF outbreaks.

In 2014, there were 14 communities identified with outbreaks of genotype III, including Baiyun, Beijing, Dadong, Guangta, Jinsha, Licheng, Liurong, Meihuacun, Nancun, Shadong, Shiweitang, Tangjing, Tongde, and Wushan. In contrast, there were only two communities with outbreaks of genotype I, i.e., Dasha and Shayuan, which were both included as the control group in subsequent analyses. Because the size of the permanent resident population and the type of local environment were similar to those of control communities, we selected Tangjing and Liurong as the study groups and compared the capacities of genotypes I and III to drive outbreaks. Our results demonstrated that genotype III showed more driving force than genotype I in dengue outbreaks as a new genotype in 2014, with an RR value of 1.61 (95% CI: 1.47– 1.76; Table 3).

**Table 3.** Comparison of new genotype invasion and previous genotypes driving dengue outbreaks in 2013 and 2014.

| Year | Genotype | Number<br>of cases | Permanent<br>resident<br>population | Incidence rate<br>(100,000) | RR | 95% CI |
| --- | --- | --- | --- | --- | --- | --- |
| 2014 | III | 1442 | 131015 | 1100.64 | 1.61 | (1.47–1.76) |
|  | I | 765 | 111963 | 683.26 | 1 | -- |
| 2013 | III | 177 | 57192 | 309.48 | 2.49 | (1.89–3.28) |
|  | I | 70 | 56329 | 124.27 | 1 | -- |

Note: RR, relative risk; 95% CI, 95% confidence interval of RR value.

In 2013, only Shiweitang was found to be a site of a genotype III outbreak, whereas four communities, including Zhuguang, Zhongnan, Kuangquan, and Jianggao, had genotype I outbreaks. Similarly, when searching for a suitable control community based on the size of the permanent resident population and the type of local environment, we selected Zhuguang to compare the capacity of genotype I for driving outbreaks with that of genotype III in Shiweitang. The results showed that the risk of genotype III as a new genotype for driving outbreaks in 2013 was 2.49 times higher than that of genotype I (95% CI: 1.89–3.28; Table 3).

### Association of different genotypes with outbreaks over the years

We used a linear regression model to analyze the relationships between the epidemic capacities of different genotypes where the community incidence caused by different genotypes was selected as the dependent variable, the factorial genotype was included as the independent variable with genotype I being the reference, and population density was entered as the adjusting variable. Both univariate and multivariate analyses demonstrated that genotype III showed a positive correlation and the greatest regression coefficient in magnitude with statistical significance (Table 4). Additionally, there was no statistically significant association between genotypes II and I.

**Table 4.** Results of linear regression analysis of genotype III in driving the outbreaks

| **Model** | **$\beta$** | ***Se*** | **$P^\beta$** | ***F*** | **$P^f$** |
| --- | --- | --- | --- | --- | --- |
| Model 1 | 393.00 | 134.5 | 0.006 | 5.50 | 0.009 |
| Model 2 | 390.20 | 138.14 | 0.008 | 3.558 | 0.025 |

Note: Model 1 is a univariate model (independent variable: genotype); model 2 is based on model 1 with the population density included as the adjusting factor; $P^\beta$ represents the *P* value of the *t* test of the regression coefficient; and $P^f$ represents the *P* value of the *F* value test of the regression equation.

### Sequence analysis of the genotype III coding region and each protein gene

In this study, genotype III first appeared in October 2013. We compared the coding sequences of the 2013 strain (KX225487) with those of the 2014–2016 strains and found that the average similarity ratio was 99.88%, indicating that base mutations occurred after the genotype invasion. Specifically, base mutations occurred in all three structural proteins and seven nonstructural proteins. The gene sequence encoding NS3 protein had the lowest average similarity ratio (99.82%), followed by the gene sequence encoding E protein (99.86%), suggesting that NS3 and E gene sequences experienced faster mutation after the DENV-1 genotype III invaded Guangzhou (Table 5, Additional File Figure S2).

**Table 5.**
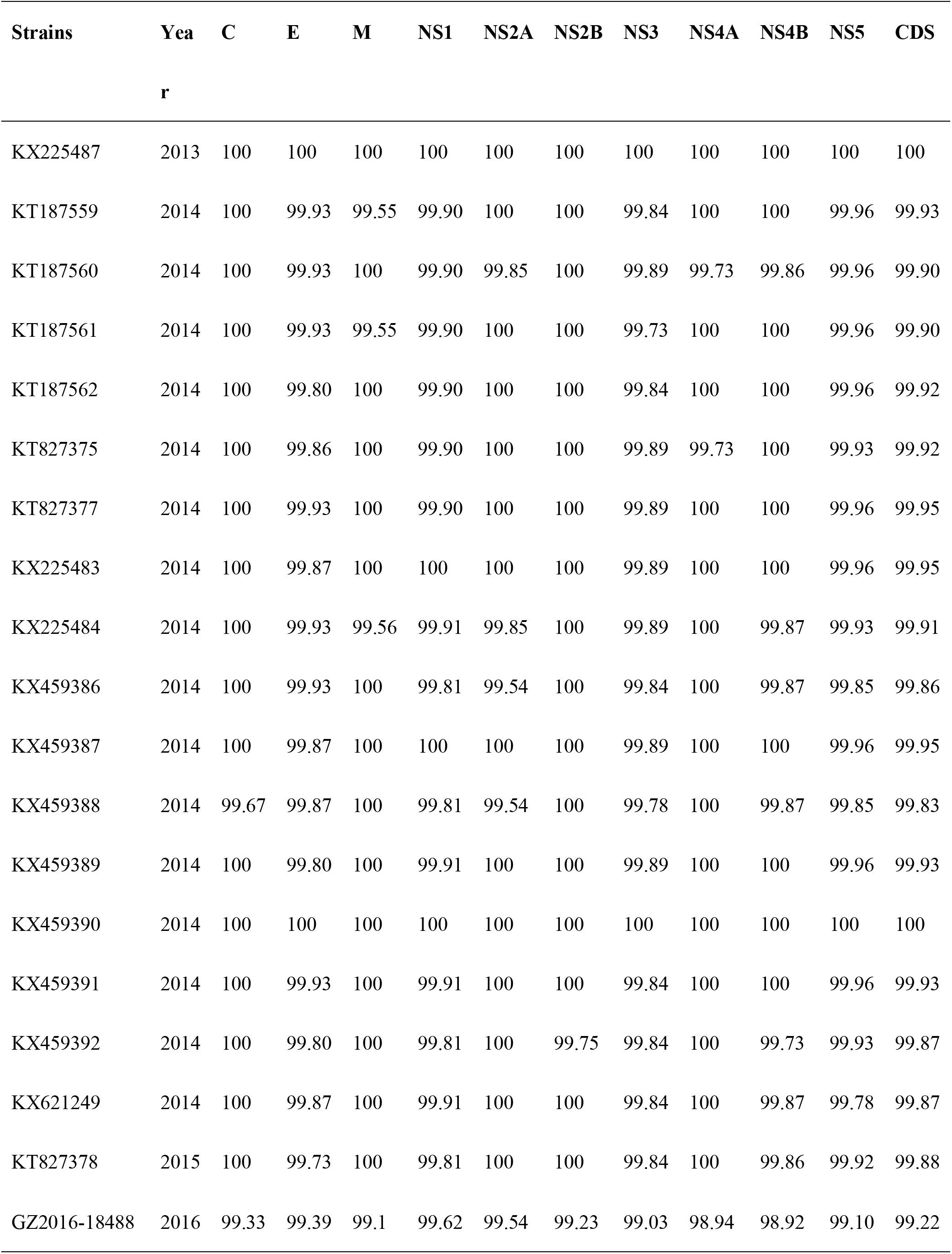
Sequence similarity between the 2013 strain (KX225487) and strains isolated in 2014–2016 (%)

The similarity ratio showed a downward trend each year, with the highest similarity ratios for coding sequences in 2013 and 2014 (average: 99.92%), followed by 2015 (99.88%) and 2016 (lowest, 99.22%). Similar trends were observed for E, NS1, NS3, and NS5 proteins when evaluating specific protein sequences. Specifically, the similarity ratios of the E gene region were 99.90% in 2014, 99.73% in 2015, and 99.39% in 2016; those in the NS1 gene region were 99.91% in 2014, 99.81% in 2015, and 99.62% in 2016; those in the NS3 gene region were 99.87% in 2014, 99.84% in 2015, and 99.03% in 2016; and those in the NS5 gene region were 99.94% in 2014, 99.92% in 2015, and 99.91% in 2016.

Gene regions encoding C, M, NS2A, NS2B, and NS4A proteins were conserved in 2014 and 2015, and the similarity ratios of most sequences reached 100%. However, the sequence similarity ratios of the 2016 strain in either coding region or each protein region (C, E, M, NS1, NS2A, NS2B, NS3, NS4A, NS4B, and NS5) decreased to varying degrees, among which the similarity ratio of the NS4B gene region was the lowest (98.92%), followed by those of the NS4A, NS3, M, NS5, NS2B, C, E, NS2A, and NS1 protein sequences at 98.94%, 99.03%, 99.10%, 99.10%, 99.23%, 99.33%, 99.39%, 99.54%, and 99.62%, respectively.

## Discussion

In this study, genotype replacement and co-circulation of multiple genotypes were observed in DENV-1 outbreaks in Guangzhou, China. We found that DENV-1 genotype II was responsible for 2002 outbreaks. However, no large-scale genotype II outbreaks were detected from viral isolate samples in the following years. Additionally, in 2006, genotype I was first identified as a new genotype invasion and was then found to co-circulate with genotype II in 2007. Similar findings were observed until 2013, when the invasion of new genotype III occurred, replacing genotype II to co-circulate with genotype I. Using complete genome sequence analysis and comparative analysis of the epidemic capacity of different genotypes, we found that there were new DENV-1 genotype invasion events in all major outbreak years in Guangzhou, and the appearance of new genotypes was highly correlated with the scale of the outbreak.

In terms of the unprecedented large-scale dengue outbreak in Guangzhou in 2014, we observed invasion of a new genotype (genotype III) of DENV-1 in addition to genotype I, which had dominated the epidemics in previous years. This high-intensity outbreak was mainly driven by the new genotype III. Phylogenetic analysis also showed that DENV-1 genotype III, isolated in 2013 and 2014, had high similarity with strains from India (JQ922548/India/2005, JQ917404/India/2009) and Singapore (KM403584/Singapore/2013).

New genotype invasion is an important feature of unusual outbreaks in major dengue-endemic regions worldwide. Various serotypes, genotypes, and their lineage clades are different in terms of viral virulence and epidemic capacity [2, 17], with outbreaks characterized by new genotype invasions being the most remarkable. Introduction of new serotypes or genotypes can often change the dominant circulating viral strains in a region. For example, the epidemics in Malaysia in 1993–1995 could be traced back to the invasion of DENV-3 genotype II in Thailand in 1962–1987. The Malaysian DENV-3 isolate used to be genotype I prior to the invasion and was soon replaced by a high-intensity outbreak of DENV-1 and DENV-2 cocirculation during 1995–1998. Changes in different serotypes repeatedly caused intense outbreaks after 2000, with DENV-1 responsible for the most serious epidemics [23–25]. In India, DENV-2 genotype V was gradually replaced by genotype IV from 1967 to 1996, which was accompanied by severe epidemics [26]. By 2003–2004, a phylogenetic analysis revealed that outbreaks were highly related to the invasion of the new DENV-3 genotype III, which eventually took over the previous DENV-2 genotype IV and became the dominant serotype and genotype [27].

Dengue-affected geographic areas are constantly expanding owing to the emergence of new genotypes. For example, DENV-3 genotype III, first identified in the Indian subcontinent, spread to Africa in the 1980s and was further disseminated to Latin America in the 1990s. Notably, the virulence of DENV-3 genotype III changed and tended to be enhanced during geographic dissemination, as demonstrated by statistically significant distribution of mild and severe cases in phylogenetic analysis [28]. In Venezuela, E gene sequence analysis of DENV-3 isolates in the 2000–2001 outbreak showed their likely origin to be a genotype III strain that had invaded from Nicaragua and Panama. This genotype continued to spread in Central America and Mexico and eventually replaced genotype V, which had been epidemic in Venezuela from the 1960s to the 1970s [29]. In Central and South America, genotype invasions occurred more frequently. DENV-3 genotype III spread twice from the Caribbean to Brazil and was introduced to Paraguay at least three times [30], causing serious dengue outbreaks in both countries and surrounding areas. Phylogenetically, Ecuadorian DENV strains were also associated with isolates with Latin American origin [31]. The outbreak of DENV-2 in Puerto Rico originated from the invasion of the new Asian genotype IIIb, and since then, the clade has been cocirculating in the country with another lineage from the Western hemisphere [32]. With regard to corresponding variations in virulence, previous studies based on the E gene suggested that positive selection occurred at several amino acid positions of the E gene, and such point mutations resulted in not only enhanced transmission but also increased viral virulence.

DENV-1 is most important serotype causing serious outbreaks in China, Southeast Asia, and the South Pacific in recent years. DENV-1 consists of five genotypes (I–V) according to previous phylogenetic analyses based on the E gene, and there are clades of varied sequence features within each genotype [17]. The strain of DENV-1 causing the outbreak in the South Pacific during 1988–1989 was only distantly phylogenetically related to the dominant strains in the region and was much closer related to the American strain, suggesting that the outbreak was caused by the invasion of a new genotype rather than a sudden outbreak of a previous epidemic strain [33]. In 2001, outbreaks of three different genotypes of DENV-1 (I–III) occurred almost at the same time in Myanmar and the South Pacific region; these outbreaks could be traced back to multiple introductions from neighboring regions of Asia [34]. Similarly, there were DENV-1 outbreaks in Hawaii and Tahiti during 2001–2002, and phylogenetic analysis showed that Hawaiian isolates actually originated in Tahiti through invasion of genotype IV [35]. In China, the 2004 DENV-1 outbreak in Zhejiang was related to an imported case of a patient who had traveled to Thailand [36]. In general, dengue outbreaks in China tend to be exclusively caused by imported cases.

Singapore experienced their largest outbreak in history from 2013 to 2014. DENV-1 replaced DENV-2 as the main serotype in circulation, resulting in a total of 40,508 cases, including 22,170 cases in 2013 and 18,338 cases in 2014 with incidence rates as high as 410.6/100,000 and 335.0/100,000, respectively. The outbreak was ultimately confirmed to be caused by the invasion of a DENV-1 genotype III variant [37]. Further analysis of the genetic variation of the new genotype during the epidemic course revealed that there were three different variants in genotypes generated during the local epidemic course in Singapore. These variants exhibited different temporal and spatial distribution patterns with regard to driving the outbreak [19]. In the same year, the largest outbreak of DF was also observed in Taiwan, with a total of 15,732 cases reported, including 136 cases of dengue hemorrhagic fever and twenty deaths, primarily caused by the new genotype of DENV-1 [38]. Thus, DENV-1 genotype III was a key factor of large-scale outbreaks in Southeast Asia, and the successive unprecedented large-scale outbreaks in Singapore and Taiwan during 2013–2014 were both closely related to the invasion of DENV-1 genotype III.

Variations in genotypes and their clades have been shown to cause severe dengue epidemics and cases [38]. In Myanmar, 15,361 cases of dengue hemorrhagic fever/dengue shock syndrome and 192 deaths were reported in 2001, and 95% of the cases were caused by DENV-1 [39]. Further phylogenetic studies have shown that the two lineages of the DENV-1 genotype I were previously unknown to the region and probably caused by new variations generated from stochastic epidemic events [40]. In 2015, DENV-4 genotype I clade C caused severe cases in southern India, and sequence analysis demonstrated that there were mutations in amino acid sites involved in viral replication and epitope presentation [41].

In this study, we demonstrated that DENV-1 new genotype invasion typically caused dengue outbreaks in Guangzhou, particularly in 2006 and 2013–2014, whereas DENV-1 genotype II was the main epidemic genotype in 2003 and before. However, after invasion of the DENV-1 genotype I in 2006, Guangzhou soon experienced the largest outbreak of genotype I, with 765 local cases reported [42]. During the middle and late stages of the epidemic in 2013, DENV-1 genotype III was introduced to Guangzhou as a new genotype invasion. As a result, 1249 local cases were reported, of which 78 cases developed into severe disease, representing the largest outbreak since 2002. DENV-1 genotype III continued to cause outbreaks in 2014 and eventually led to a record-breaking outbreak, with a total of 37,340 local cases reported. This number was over 2.4 times the sum of all cases reported from 1978 to 2013, and 14,000 cases of hospitalization, 308 severe cases, and five deaths were observed. Complete genome sequence analysis of DENV-1 showed that there were two genotypes (I and III) cocirculating in 2014. No significant variations in genotype I were observed. Therefore, this large-scale outbreak was highly associated with genotype III, and the capacity of genotype III for driving outbreaks was stronger than the capacities of genotypes I and II. DENV-1 genotype III strains appeared only in April 2015 and re-emerged in 2016. Moreover, studies have shown that secondary infections played a negligible role in severe cases during the 2014 outbreak [43], suggesting that new genotypes could increase the risk of developing into severe cases.

Because asymptomatic patients and patients with mild disease usually do not seek medical treatment and the patients may visit the hospital at later stages of the disease, specimens in this study were mainly from symptomatic patients, and no asymptomatic individuals (and few patients with mild disease) underwent virus isolation. Therefore, the studied strains may not have represented the entire infected population. In addition, owing to financial constraints, the number of isolated strains and self-sequenced complete genome data in our study were still limited.

Our current findings demonstrated that serotype replacement and variations in genotypes and clades, particularly new genotype invasion, were important features of large-scale outbreaks of DF. However, future prospective epidemiological and phylogenetic studies are required to further clarify the genetic variations of new genotypes with different genotypes and clades co-circulating in affected areas as well as the epidemic capacities and scales of resulted outbreaks. Moreover, additionally epidemiological studies of severe cases are needed to comprehensively evaluate new genotype invasion and its capacity for driving local epidemics and causing severe disease. Such studies could provide valuable scientific support for prevention and control efforts as well as early detection in dengue-affected areas.

## Acknowledgements

We would like to thank all 12 district CDCs in Guangzhou and their staff for participating in this study. We would also like to thank Fuchun Zhang from Guangzhou 8^th^ People’s Hospital and Xiaohong Zhou and Xiaoguang Chen from Southern Medical University in China for valuable suggestions.

## Consent for publication

All authors approved the final version of this manuscript.

## Author contributions

QLJ, JHL, and ZCY generated the idea and organized the study; QLJ, SW, ZJH, LHY, and MMM drafted and modified the manuscript. QLJ, ZJH, and MMM conducted field surveys and collected data; QLJ, LYJ, and ZJB conducted the laboratory test; QLJ, SW, ZJH, and JM conducted data analysis.

## Supporting information

**Additional File 1: Table S1. Specific primers for amplification and sequencing of dengue virus serotype 1.** Total 35 pairs of primers were designed for sequencing DENV-1 genome.

**Additional File 2: Table S2. Distribution of DENV-1 genome sequences in different countries and regions.** Total 1679 DENV-1 complete genome sequences were analyzed, including 97 strains from China and 1582 strains from other countries and regions.

**Additional File 3: Fig S1. DENV-1 complete genome sequence phylogenetic tree.** 1631 genome sequences from GenBank and 48 genome sequences from research team were aligned using Mafft software (version 7). Phylogenetic tree was constructed with the neighbor-joining method with Kimura 2-parameter corrections of multiple substitutions using Mega software (version 7.0). Virus strains from China are indicated by read lines.

**Additional File 4: Fig S2. Phylogenetic trees for each protein and complete coding sequence of DENV-1 genotype III in Guangzhou**. One strain in 2013 and 18 strains from 2014 to 2016 of DENV-1 genotype III from Guangzhou were were aligned using Mafft software (version 7). Phylogenetic tree was constructed with the UPGMA method using Mega software (version 7.0).

